# Microbial community-based bacterial protein from soybean-processing wastewater as a value-added alternative fish feed ingredient for Asian sea bass (*Lates calcarifer*)

**DOI:** 10.1101/2023.01.18.524524

**Authors:** Ezequiel Santillan, Fanny Yasumaru, Ramanujam Srinivasan Vethathirri, Sara Swa Thi, Hui Yi Hoon, Diana Chan Pek Sian, Stefan Wuertz

## Abstract

As the global demand for food increases, aquaculture plays a key role as the fastest growing animal protein sector. However, existing aquafeeds contain protein ingredients that are not sustainable under current production systems. We evaluated the use of microbial community-based single cell protein (SCP), produced from soybean processing wastewater, as a partial fishmeal protein substitute in juvenile Asian seabass (*Lates calcarifer*). A 24-day feeding trial was conducted with a control fishmeal diet and a 50% fishmeal replacement with microbial community-based SCP as an experimental group, in triplicate tanks containing 20 fish each. Both diets met the protein, essential amino acids (except for lysine), and fat requirements for juvenile Asian sea bass. The microbial composition of the SCP was dominated by the genera *Acidipropionibacterium* and *Propioniciclava*, which have potential as probiotics and producers of valuable metabolites. The growth performance in terms of percent weight gain, feed conversion ratio (FCR), specific growth rate (SGR), and survival were not significantly different between groups after 24 days. The experimental group had less variability in terms of weight gain and FCR than the control group. Overall, microbial community-based protein produced from soybean processing wastewater has potential as a value-added feed ingredient for sustainable aquaculture feeds.

## 1. Introduction

It is estimated that 1,250 million tons of animal-derived protein would need to be produced per year in 2050 to meet the global demand at current consumption levels (Ritala et al., 2017). However, increasing meat production for this purpose is not sustainable because the conversion of plant to meat protein is inefficient (Linder, 2019). Protein production by conventional agriculture-based food supply chains is also a major issue in terms of global environmental pollution. The livestock sector produces 14.5% of the global anthropogenic greenhouse gas (GHG) emissions, and about 33% of the land surface and 75% of freshwater resources are used for livestock or crop production (EC, 2020). Thus, new solutions for protein production are needed to ensure sustainable food security.

Protein can be produced through the cultivation of various microbes, which is generally denoted as microbial protein or single cell protein (SCP) (Anupama & Ravindra, 2000). A recent life cycle assessment (LCA) showed a lower environmental impact of SCP production compared to that of soybean meal manufacturing (Spiller et al., 2020). SCP grown on food processing wastewaters (FPWW) holds promise as a sustainable source of protein in animal feed (Vethathirri et al., 2021). Collectively, FPWW possess attractive features for SCP production, such as a continuous global production of process water rich in dissolved carbon (C), nitrogen (N) and phosphorus (P) compounds (Muys et al., 2020). These wastewaters are also considered to be free of pathogens, heavy metals, and other harmful contaminants. FPWW can support microbial growth in bioreactors, from which cells can be recovered and dried leading to products suitable for animal feed ingredients, while concurringly avoiding the cost of treating such wastewaters, and hence representing an important step toward a circular bioeconomy (Durkin et al., 2022).

The supply of seafood, wild-caught and farmed, for human consumption was around 175 million tons in 2020, and is the largest protein industry worldwide (Boyd et al., 2022; Jones et al., 2020). Aquaculture accounted for about 50% of that sector and 21% of the total animal protein produced (FAO-UN, 2020). As the global population is predicted to increase by 30% in 2050 compared to 2018, it is estimated that aquaculture production would need to increase by 57.2% to meet the growing demand for seafood (Boyd et al., 2022). GHG emissions by aquaculture are estimated as 0.49% of global emissions (Boyd & McNevin, 2015; MacLeod et al., 2020). Nevertheless, high trophic level aquaculture species still rely on fishmeal and fish oil derived from wild capture fisheries (Naylor et al., 2009; Slater et al., 2018; Tacon et al., 2022). SCP represents a promising alternative to fishmeal as a protein source in aquaculture (Jones et al., 2020), as it does not compete with the human food supply. In fact, it can transform the so-called “waste” from the human food industry into a valuable, nutritious, and sustainable animal feed ingredient. Thus, it may offer solutions in aquaculture across a wide variety of products and production approaches, but considerable research, development, and scale-up efforts are required to reach its potential (Carter & Codabaccus, 2022; Verstraete et al., 2022).

Traditionally, SCP production involves the use of single strains in axenic cultures, incurring high initial and operational costs. For example, a recent study used pure and enriched cultures of purple bacteria as a protein ingredient (5-11% protein replacement) in whiteleg shrimp nursery feed (Alloul et al., 2021b). An alternative would be a whole-community approach for SCP production, leveraging the microorganisms already present in the wastewater (Vethathirri et al., 2021), but remains largely unexplored. A recent study showed that microbial community-based SCP produced directly from soybean processing wastewaters contained essential amino acids for fish as well as taxa with probiotic potential (Vethathirri et al., 2022). However, the feasibility of using such SCP as fish feed ingredient for aquaculture has yet to be established. Hence, the aim of this study was to evaluate the use of microbial community-based SCP, produced from soybean processing wastewater, as a value-added alternative ingredient to fishmeal for juvenile Asian seabass (*Lates calcarifer*) aquaculture.

## 2. Materials and methods

### 2.1 Fish source and experimental culture conditions

Unvaccinated Asian sea bass (*Lates calcarifer*, also known as barramundi) juveniles were obtained from a commercial hatchery (Allegro Aqua, Singapore). Fish were fed with a commercial sea bass diet (43% protein, Uni-President, Vietnam) for a period of three weeks to acclimate the fish to lab conditions prior to the start of the 24-d feeding trial. Juvenile sea bass of 19.7 ± 2.6 g were stocked in rectangular glass aquaria (0.58 m * 0.58 m * 0.38 m, 100-L water volume) at a stocking density of 20 fish aquaria^-1^ (m = 20), with three aquaria allotted per dietary treatment (n = 3). The glass aquaria were connected to a recirculating system with mechanical and biological filtration and aeration was supplied to each aquarium. Water temperature ranged from 27 to 29 °C. Other water quality parameters, *i*.*e*., dissolved oxygen (> 4 mg L^-1^), pH (7.0 - 7.9), salinity (15 ppt), ammonia (< 0.5 ppm) and nitrite (< 0.5ppm) were kept within acceptable levels for the species. The study was approved by the Institutional Animal Care and Use Committee (IACUC) under number 202003-155A.

### 2.2 Experimental diets and feeding protocol

A control fishmeal-based (FM) diet and an experimental 50% fishmeal replacement diet with microbial community-based SCP (50SCP), were formulated to contain at least 45% protein and 12% fat (Table 1). These were prepared with 30% fishmeal (FM) and with 50% replacement of the fishmeal with SCP (50SCP), while adjusting the other ingredients to maintain the crude protein and fat levels (Table 1). The nutritional composition of both diets is detailed in Table 2. Diet preparation involved mixing the dry feed ingredients first, followed by the addition of the liquids, including 20% v/v water, in a stand mixer (Model 5 QT, KitchenAid, St Joseph, MI, USA). The resulting mash was passed through a meat grinder attached to the stand mixer and fitted with a 3-mm diameter die. The moist strings were dried overnight in a ventilated drying oven at 45 °C. The dry strings were hand-broken and stored in air-tight plastic containers at 4 °C until used. Each diet was randomly assigned to three replicate glass tanks. Both diets were fed twice a day (0900 hours, 1600 hours) to apparent satiation, and feed intake was measured daily. Fish were group-weighed at the beginning and individually at the end of the 24-day feeding trial.

**Table 1.** Formulation of the fishmeal (FM) control diet and microbial community-based single-cell protein experimental diet (50SCP – 50% fishmeal replacement) used in the feeding trial with Asian sea bass (*Lates calcarifer*).

| **Ingredient (% w/w)** | **FM** | **50SCP** |
| --- | --- | --- |
| Fishmeal | 30.0 | 15.0 |
| Microbial community-based single-cell protein (SCP) | 0.0 | 15.0 |
| Poultry by-product meal | 13.0 | 16.0 |
| Squid liver meal | 10.0 | 10.0 |
| Dried yeast | 3.0 | 3.0 |
| Fish hydrolysate | 5.0 | 5.0 |
| Blood meal | 4.0 | 4.0 |
| Wheat flour | 18.5 | 15.5 |
| Wheat gluten | 5.0 | 5.0 |
| Soy lecithin | 2.5 | 2.5 |
| Fish oil | 8.0 | 8.0 |
| Vitamin premix | 0.2 | 0.2 |
| Mineral premix | 0.2 | 0.2 |
| Taurine | 0.5 | 0.5 |
| Ascorbic 2-phosphate 35% | 0.2 | 0.2 |

**Table 2.**
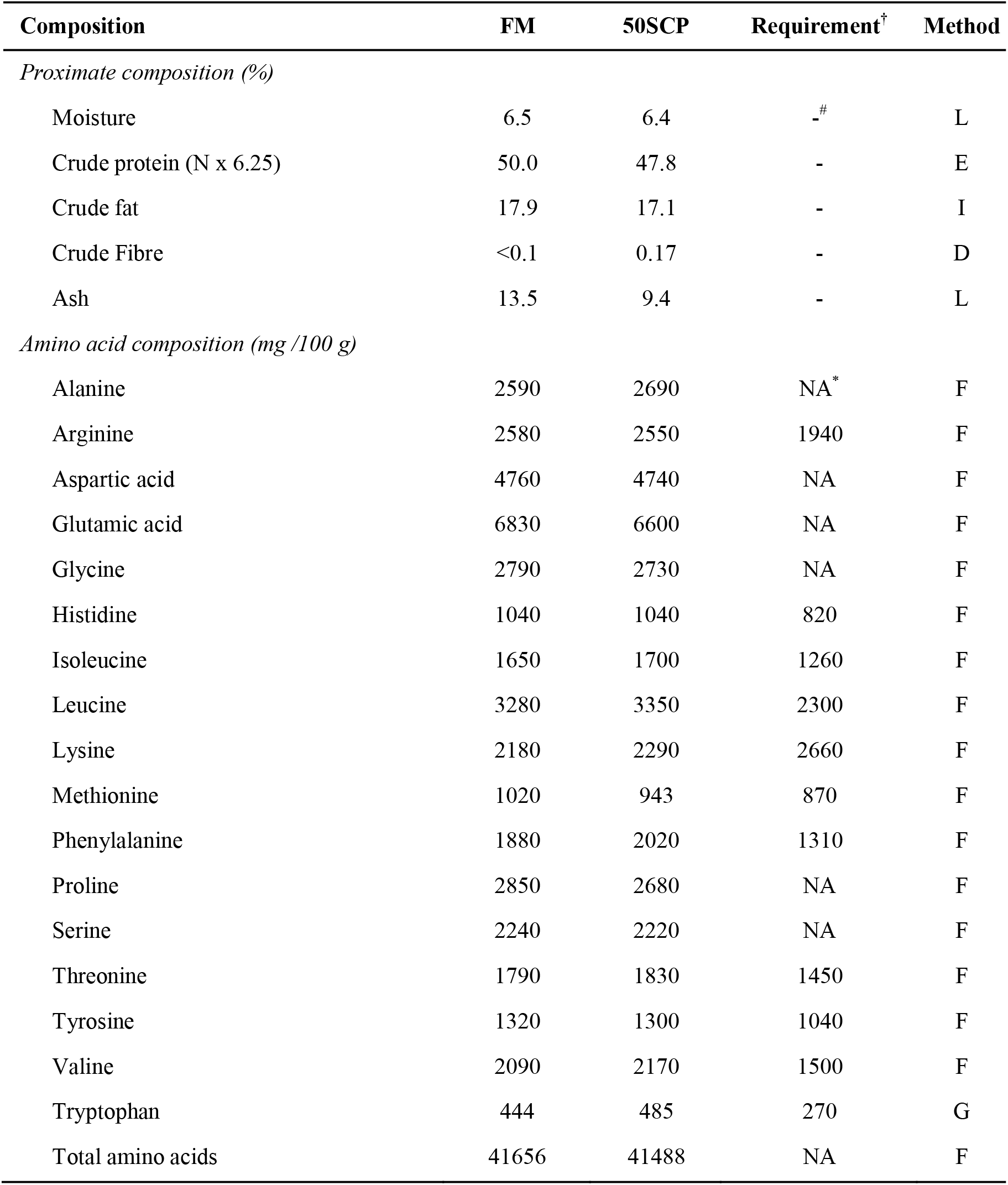
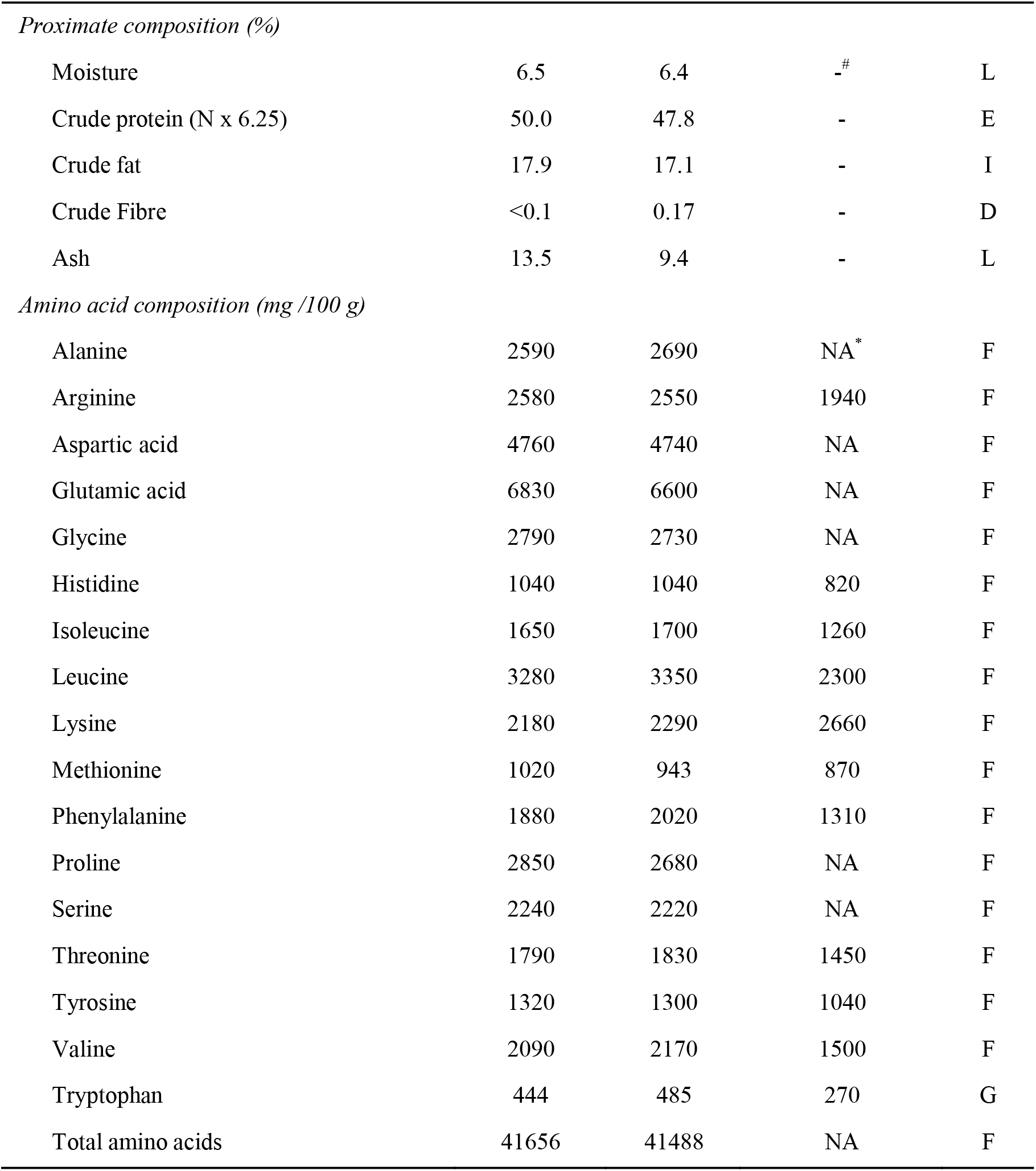
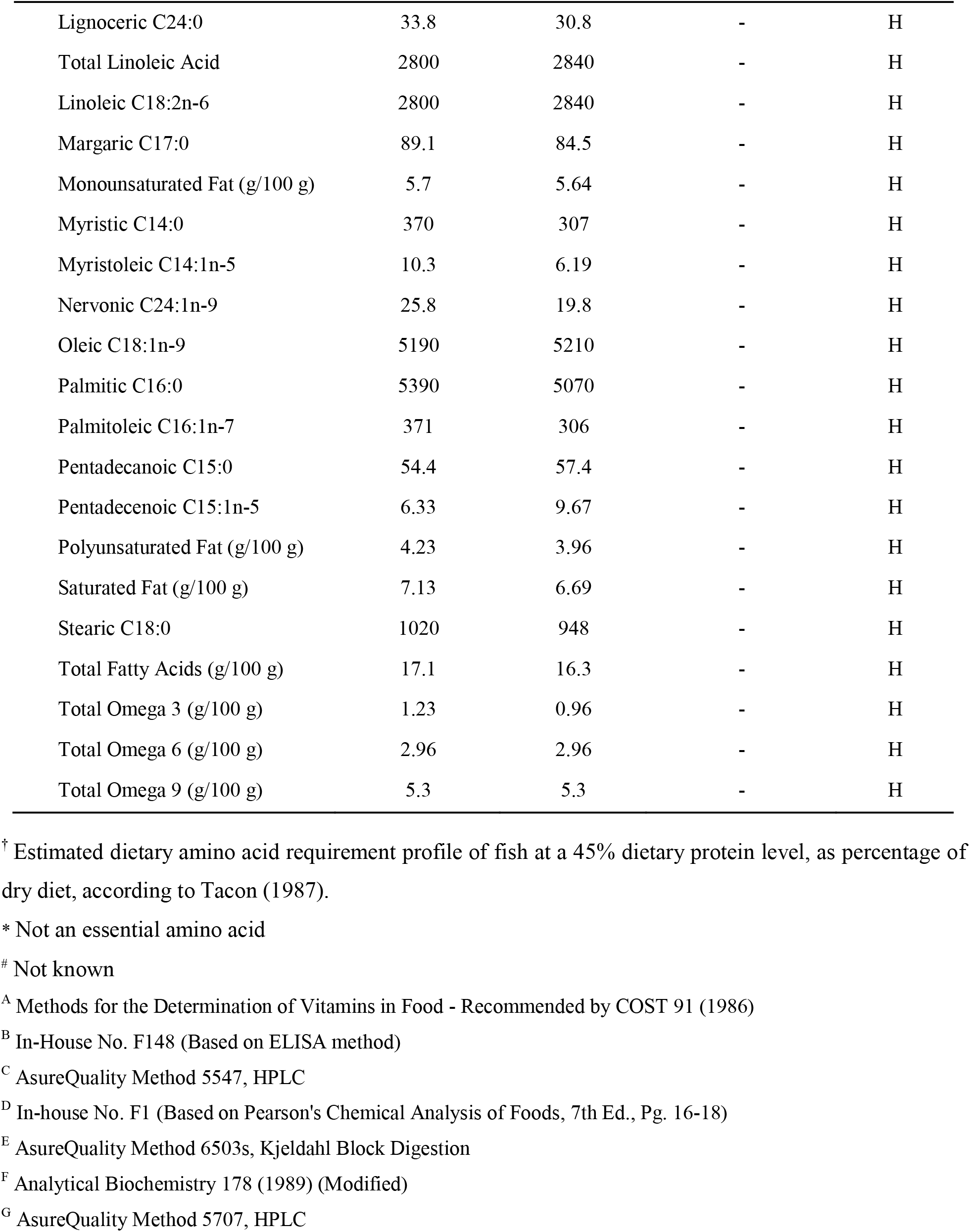
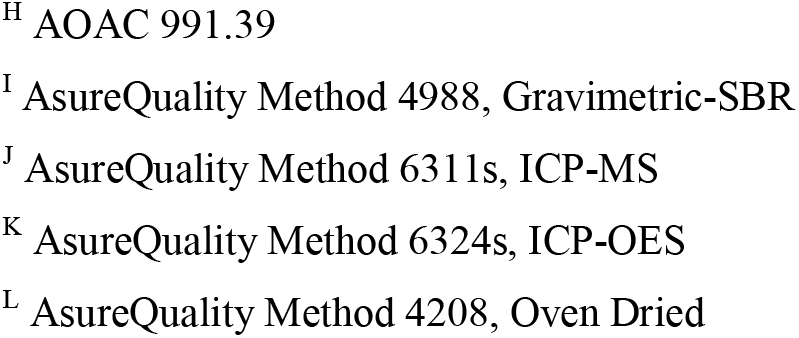
Nutritional composition (“as is” basis) of the fishmeal (FM) control diet and the experimental diet with 50% fishmeal replacement by microbial community-based single-cell protein (50SCP) used in the feeding trial with Asian sea bass (*Lates calcarifer*).

### 2.3 Bioreactor setup and operational parameters for SCP production

To produce microbial SCP, four 4-L bioreactors were operated as sequencing batch reactors (SBRs) on continuous 12-h cycles with intermittent aeration for 136 days. Reactors received wastewater from a soybean processing company in Singapore as described in Vethathirri et al. (2022). Water temperature was maintained at 30 °C, and the sludge was continuously mixed at 375 rpm. The feeding phase occurred during the initial 5-10 min of a cycle, followed by 180 min anoxic/anaerobic and 540 min aerobic phases, after which the biomass was left to settle for 60 min and 1.35 L of the supernatant was discarded. Thereafter, the reactor was filled with the same volume of soybean wastewater, starting a new cycle. The pH was measured between 6.0 – 8.5, and the dissolved oxygen (DO) concentration was controlled between 0.2 – 0.5 mg/L during the aerobic phase. Each of the SBRs employed in this study was equipped with a magnetic stir plate to ensure mixed liquor homogeneity, a pair of EasySense pH and DO probes with their corresponding transmitters (Mettler Toledo), dedicated air and feed pumps, a solenoid valve for supernatant discharge, and a surrounding water jacket connected to a re-circulating water heater. The different portions of the cycle were controlled by computer software specifically designed for these reactors (VentureMerger, Singapore).

### 2.4 Chemical analyses of fish diets and tissue

The nutritional composition, *i*.*e*., proximate composition (moisture, crude protein, crude fat, crude fibre, ash), amino acids profile, fatty acids profile, calcium, phosphorus, selenium, zinc, iron, and vitamins A, D, and B12 content of the control and experimental diets, as well as fish tissue (one sample per group) were analyzed according to standard methods (AOAC, 1995) by BV-AQ laboratories (Singapore).

### 2.5 Microbial characterization of microbial community-based SCP

Three samples of 0.5 g each were collected randomly from a 2-L bottle containing 222 g of mixed-culture SCP to be used in the aquaculture trial. These were subjected to DNA extraction and as previously described (Santillan et al., 2019). Amplicon preparation was done by PCR using the primer set 341f/785r targeted the V3-V4 variable regions of the 16S rRNA gene (Thijs et al., 2017), as described in (Santillan & Wuertz, 2022). Bacterial 16S rRNA amplicon sequencing was done in two steps as described in Santillan et al. (2020b). The libraries were sequenced on an Illumina MiSeq platform (v.3) with 20% PhiX spike-in and at a read-length of 300 bp paired-end. Sequenced sample libraries were processed following the DADA2 bioinformatics pipeline (Callahan et al., 2016). DADA2 allows inference of exact amplicon sequence variants (ASVs) providing several benefits over traditional clustering methods (Callahan et al., 2017). Illumina sequencing adaptors and PCR primers were trimmed prior to quality filtering. Sequences were truncated after 280 and 255 nucleotides for forward and reverse reads, respectively, the length at which average quality dropped below a Phred score of 20. After truncation, reads with expected error rates higher than 3 and 5 for forward and reverse reads, respectively, were removed. After filtering, error rate learning, ASV inference and denoising, reads were merged with a minimum overlap of 20 bp. Chimeric sequences (0.03% on average) were identified and removed. For a total of 3 samples, on average 26426 reads were kept per sample after processing, representing 52% of the average input reads. Taxonomy was assigned using the SILVA database (v.138) (Glockner et al., 2017).

### 2.6 Growth performance and statistical analyses

The individual growth performance of the fish was evaluated by calculating the mean weight gain (mean final weight – mean initial weight), percent weight gain (mean weight gain * mean initial weight^-1^ *100), specific growth rate (SGR = (ln_mean final weight_ – ln_mean initial weight_) * trial duration^-1^ *100), feed conversion ratio (FCR = mean feed intake * mean weight gain^-1^), and survival rate. Univariate analysis of growth performance parameters were done via Welch’s t-test assuming unequal variances in R (v.3.6.3). All reported P-values for statistical tests in this study were corrected for multiple comparisons using a false-discovery rate (FDR) of 5% (Benjamini & Hochberg, 1995). Heat maps for bacterial relative abundances of the SCP were constructed using the DivComAnalyses package (v.0.9) in R (Constancias & Sneha-Sundar, 2022).

## 3. Results and discussion

Both the control fishmeal-based diet (FM) and the microbial community-derived SCP experimental diet (SCP50) met the protein (>45%) and fat (>12%) requirements for juvenile Asian sea bass (Table 1). They were also formulated to provide essential amino acids to fish; however, at 18% and 14% in the FM control diet and 50SCP experimental diet, respectively, lysine was below the required level (Table 2). The remaining essential amino acid requirements were met, in agreement with a recent study by Vethathirri et al. (2022) where microbial community-derived SCP biomass produced from soy bean FPWW in lab-scale reactors was reported to fully or partially meet the essential amino acids requirements for fish and shrimp.

No significant differences in growth performance were observed between fish fed with the FM control and those that received the experimental microbial community-based 50SCP diet (Table 3). The initial weight of the fish in the group fed with the 50SCP experimental diet was significantly lower (Welch’s t-test P_adj_ < 0.001) than the control, despite the initial random assignment, but no significant difference was observed in the weight gain at the end of the feeding trial (Welch’s t-test P_adj_ = 0.30) (Table 3, Figure 1). The feed intake of the group fed with the FM control diet was not significantly different from that of the group fed with the 50SCP mix experimental diet. Hence, no significant difference was observed for the feed conversion ratio (FCR) and specific growth rate (SGR). No mortality was observed during the feeding trial in both groups (Table 3). Furthermore, the group fed with FM control diet presented a 4.4- and 10.3-fold greater variability in percent weight gain and FCR than the group fed with the 50SCP experimental diet, respectively, suggesting that an SCP replacement diet could also lead to more homogeneous growth of farmed Asian sea bass.

**Table 3.** Growth of juvenile Asian sea bass (*Lates calcarifer*) fed with a fishmeal (FM) control diet or an experimental diet with 50% fishmeal replacement with microbial community-based SCP (50SCP) over 24 d. Values are means ± s.d.m. (n = 3).

| Parameter | FM | 50SCP | P<sub>adj</sub><sup>†</sup> |
| --- | --- | --- | --- |
| Initial weight (g) | 22.1 ± 0.21 | 17.4 ± 0.14 | < 0.001 |
| Final weight (g) | 38.6 ± 5.14 | 31.6 ± 0.78 | 0.28 |
| Weight gain (g) | 16.5 ± 5.30 | 14.3 ± 0.88 | 0.69 |
| Weight gain (%) | 74.8 ± 24.7 | 82.1 ± 5.58 | 0.69 |
| Feed intake (g) | 19.8 ± 1.56 | 16.8 ± 0.73 | 0.15 |
| Feed conversion ratio<sup>§</sup> | 1.26 ± 0.31 | 1.18 ± 0.03 | 0.69 |
| Specific growth rate (% day<sup>-1</sup>)<sup>‡</sup> | 2.30 ± 0.57 | 2.50 ± 0.13 | 0.69 |
| Survival (%) | 100 | 100 | - |

<sup>†</sup> Welch's t-test P-values adjusted at 5% FDR
<sup>§</sup> FCR = mean feed intake \* mean weight gain<sup>-1</sup>
<sup>‡</sup> SGR = (ln<sub>mean final W</sub> – ln<sub>mean initial W</sub>) \* trial duration<sup>-1</sup> \* 100

**Figure 1.**
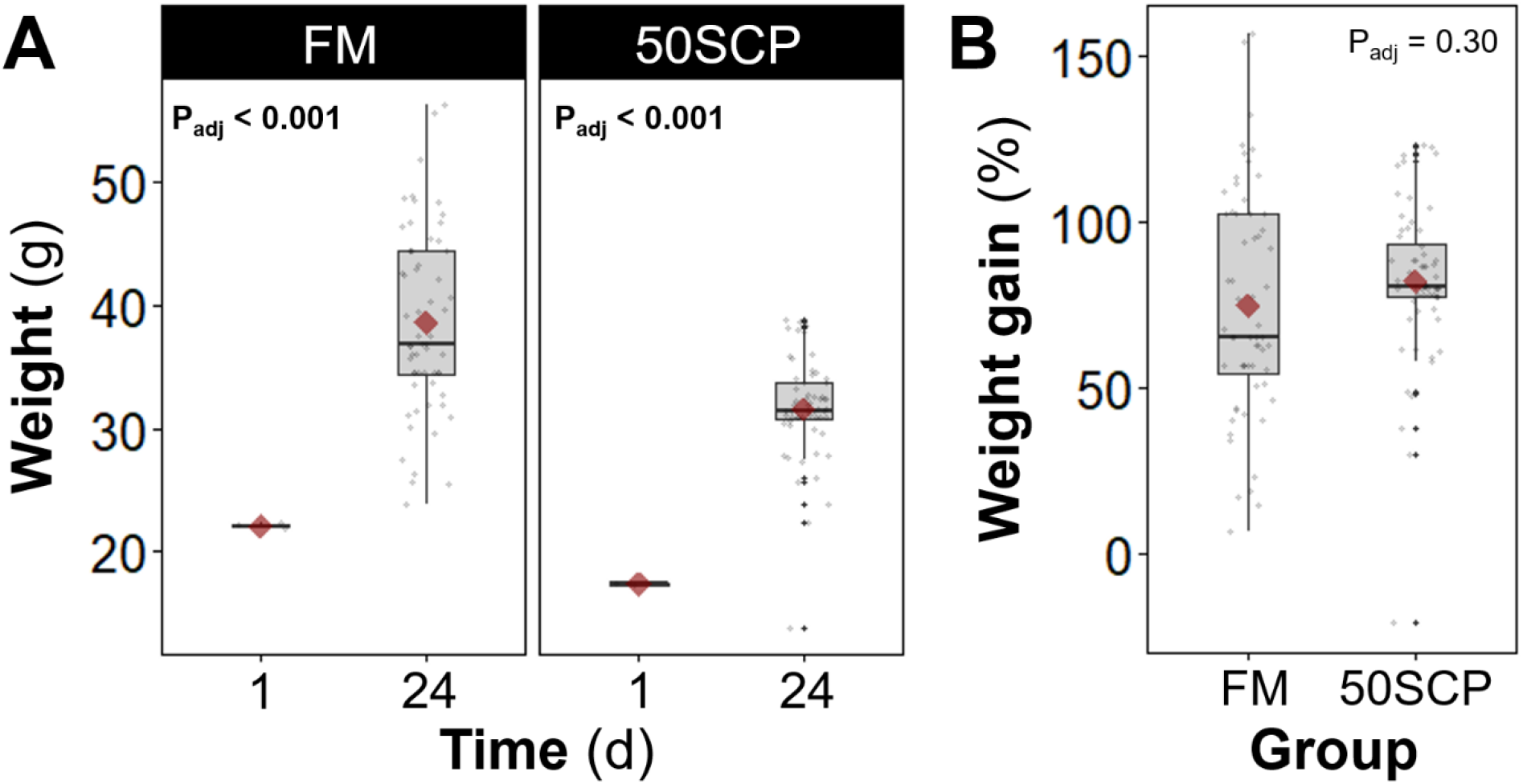
Weight gain of juvenile Asian sea bass (*Lates calcarifer*) fed with a fishmeal-based (FM) control diet or an experimental diet with 50% fishmeal replacement with microbial community-based SCP (50SCP) over 24 d. Panels represent (**A**) initial and final fish weight, and (**B**) percent weight gain per group. The box bounds the interquartile range (IQR) divided by the median, and Tukey-style whiskers extend to a maximum of 1.5 times the IQR beyond the box. Red diamonds display mean values. Grey dots represent values for each individual fish (m = 20) in each of three replicate tanks per group on d24. Welch’s t-test P-values adjusted at 5% FDR shown within panels. Initial values (d1) are averages for each tank, calculated by simultaneously weighing all the fish in each tank and dividing by 20.

To our knowledge, ours is among the first fish feeding trials with microbial community-derived SCP in the feed at a relatively high fishmeal replacement level (50%) with microbial protein. Other studies have investigated the use of pure or enriched cultures of purple non-sulfur bacteria (PNSB) as aquafeed additives for the whiteleg shrimp (*Litopenaeus vannamei*) at protein replacement levels of 1.2-5.8% (Chumpol et al., 2019) or 5–11% (Alloul et al., 2021b), and for barramundi (*Lates calcarifer*) at 33-100% protein replacement levels (Delamare-Deboutteville et al., 2019). These studies reported growth performance for different protein replacement levels that was comparable to that of commercial fishmeal (Alloul et al., 2021b; Delamare-Deboutteville et al., 2019), as well as probiotic properties against *Vibrio* pathogens (Alloul et al., 2021b; Chumpol et al., 2019). In our study, the nutritional composition of the fish tissue at the end of the trial was similar for the groups fed either FM control or 50SCP experimental diets in terms of crude protein, fat, most amino acids and fatty acids, as well as some vitamins and minerals (Table 4). Nevertheless, lower concentrations were observed in the fish tissue of the 50SCP experimental group for lysine (−17.6%), vitamin A (−68.8%), DHA (−20.7%), EPA (− 16.4%), and total n-3 fatty acids (−16.5%) than in the FM control group.

**Table 4.** Nutritional composition (“freeze-dried” basis) of juvenile Asian sea bass (*Lates calcarifer*) fish tissue fed with fishmeal (FM) control diet or a fishmeal replacement diet containing 50% microbial community-based SCP (50SCP) over 24 d.

| Composition | FM | 50SCP | Method |
| --- | --- | --- | --- |
| *Proximate composition (%)* |  |  |  |
| Moisture | 7.5 | 8.4 | L |
| Crude protein | 56.0 | 55.0 | E |
| Crude fat | 23.0 | 24.1 | I |
| Crude Fibre | <0.1 | <0.1 | D |
| *Amino acid composition (mg/100g)* |  |  |  |
| Alanine | 3560 | 3530 | F |
| Arginine | 3290 | 3230 | F |
| Aspartic acid | 5900 | 6480 | F |
| Glutamic acid | 6880 | 6670 | F |
| Glycine | 3890 | 3810 | F |
| Histidine | 1120 | 1140 | F |
| Isoleucine | 2030 | 1900 | F |
| Leucine | 3730 | 3510 | F |
| Lysine | 4090 | 3370 | F |
| Methionine | 1470 | 1410 | F |
| Phenylalanine | 2060 | 1990 | F |
| Proline | 2470 | 2470 | F |
| Serine | 2100 | 2000 | F |
| Threonine | 2180 | 2080 | F |
| Tyrosine | 1420 | 1350 | F |
| Valine | 2250 | 2180 | F |
| Tryptophan | 458 | 417 | G |
| Total amino acids | 49570 | 48260 | F |
| *Vitamin composition* |  |  |  |
| Vitamin A (mg kg<sup>-1</sup>) | 0.32 | 0.10 | A |
| Vitamin D3 (IU/kg) | <20 | <20 | C |
| Vitamin B12 (µg kg<sup>-1</sup>) | 0.19 | 0.16 | B |
| *Mineral composition* |  |  |  |
| Selenium (µg/100 g) | 69.6 | 65.5 | J |
| Calcium (mg/100 g) | 3640 | 3650 | K |
| Iron (mg/100 g) | 4.34 | 4.37 | K |
| Phosphorus (mg/100 g) | 1040 | 1000 | K |
| Zinc (mg/100g) | 5.24 | 5.23 | K |
| *Fatty acid composition (mg/100 g)* |  |  |  |
| Alpha Linolenic C18:3n-3 | 312 | 308 | H |
| Arachidic C20:0 | 75.3 | 74.8 | H |
| Arachidonic C20:4n-6 | 199 | 167 | H |
| Docosanoic C22:0 | 31.6 | 32.2 | H |
| Butyric C4:0 | <4 | <4 | H |
| Capric C10:0 | <4 | <4 | H |
| Caproic C6:0 | <4 | <4 | H |
| Caprylic C8:0 | <4 | <4 | H |
| Dihomo-gamma-linolenic C20:3n-6 | 49.2 | 59.5 | H |
| Docosahexaenoic C22:6n-3 (DHA) | 1180 | 936 | H |
| Eicosadienoic C20:2n-6 | 12.3 | 10.2 | H |
| Eicosapentanaeic C20:5n-3 (EPA) | 439 | 367 | H |
| Eicosatrienoic C20:3n-3 | 13.2 | 11.5 | H |
| Eicosenoic C20:1 (total) | 131 | 130 | H |
| Eicosenoic(Gondoic) C20:1n-9 | 131 | 130 | H |
| Erucic Acid C22:1n-9 | 21.4 | 16.7 | H |
| Gamma Linolenic C18:3n-6 (GLA) | 9.86 | 8.61 | H |
| Heptadecenoic C17:1n-7 | 44.8 | 46.9 | H |
| Lauric C12:0 | 14.2 | 14.7 | H |
| Lignoceric C24:0 | 27.9 | 27.9 | H |
| Linoleic C18:2n-6 | 3690 | 3790 | H |
| Margaric C17:0 | 103 | 101 | H |
| Monounsaturated Fat (g/100 g) | 8.17 | 8.92 | H |

**Table 4 – Continued**
| Composition | FM | 50SCP | Method |
| --- | --- | --- | --- |
| Myristic C14:0 | 565 | 570 | H |
| Myristoleic C14:1n-5 | 6.96 | 6.96 | H |
| Nervonic C24:1n-9 | 39.3 | 35.1 | H |
| Oleic C18:1n-9 | 7200 | 7930 | H |
| Palmitic C16:0 | 5630 | 6080 | H |
| Palmitoleic C16:1n-7 | 713 | 735 | H |
| Pentadecanoic C15:0 | 80.7 | 81.8 | H |
| Pentadecenoic C15:1n-5 | 14.7 | 17.6 | H |
| Poly Unsaturated Fat (g/100 g) | 5.97 | 5.76 | H |
| Saturated Fat (g/100 g) | 7.78 | 8.30 | H |
| Stearic C18:0 | 1210 | 1270 | H |
| Total Fatty Acid (g/100 g) | 22.0 | 23.0 | H |
| Total Omega 3 (g/100 g) | 1.94 | 1.62 | H |
| Total Omega 6 (g/100 g) | 3.95 | 4.03 | H |
| Total Omega 9 (g/100 g) | 7.39 | 8.12 | H |

<sup>A</sup> Methods for the Determination of Vitamins in Food - Recommended by COST 91 (1986)
<sup>B</sup> In-House No. F148 (Based on ELISA method)
<sup>C</sup> AsureQuality Method 5547, HPLC
<sup>D</sup> In-house No. F1 (Based on Pearson's Chemical Analysis of Foods, 7th Ed., Pg. 16-18)
<sup>E</sup> AsureQuality Method 6503s, Kjeldahl Block Digestion
<sup>F</sup> Analytical Biochemistry 178 (1989) (Modified)
<sup>G</sup> AsureQuality Method 5707, HPLC
<sup>H</sup> AOAC 991.39
<sup>I</sup> AsureQuality Method 4988, Gravimetric-SBR

A microbial community-based approach to build SCP has several potential advantages over axenic culturing (Vethathirri et al., 2021), such as a higher protein content due to synergistic interactions between different SCP-producing microorganisms and the utilization of various carbon and nitrogen sources available in the substrate (Alloul et al., 2021a; Vethathirri et al., 2022), the accumulation of intracellular components (Li et al., 2022), and process stability in terms of resistance and resilience to disturbances (Santillan et al., 2020a; Santillan et al., 2020b; Santillan et al., 2021). The microbial composition of the microbial community-based SCP employed in the experimental diet was dominated by the genera *Acidipropionibacterium* and *Propioniciclava* (Figure 2), which have potential as probiotics and producers of valuable metabolites. Taxa within the *Acidipropionibacterium* genus (previously named *Propionibacterium*) have a broad range of applications in the pharmaceutical and food industries due to their ability to produce vitamin B12, bacteriocins, and trehalose (Piwowarek et al., 2018). Indeed, there was 12 times more vitamin B12 in the experimental diet compared to the control (Table 2). Further, the species *Acidipropionibacterium acidipropionici*, which was detected in the 50SCP experimental diet, is used in the food industry as either a bio-preservative or probiotic due to its generally-recognized-as-safe (GRAS) status (Deptula et al., 2019). Some of these potential properties associated with a subset of the microbial community-based SCP could have contributed to the observed more homogeneous growth performance, in terms of FCR and weight gain, of the fish fed with the experimental diet compared to those fed with the control diet (Table 3). For example, Alloul et al. (2021b) hypothesized that PNSB would be able to colonize the gastrointestinal tract and assist the digestion of white leg shrimp given that, similar to our study, the microbial protein used was freeze-dried, which allows for bacteria to be revived. As the current knowledge of microbial community-based SCP as protein replacement in fish feed is limited, more research is needed to understand the underlying mechanisms behind our observations.

**Figure 2.**
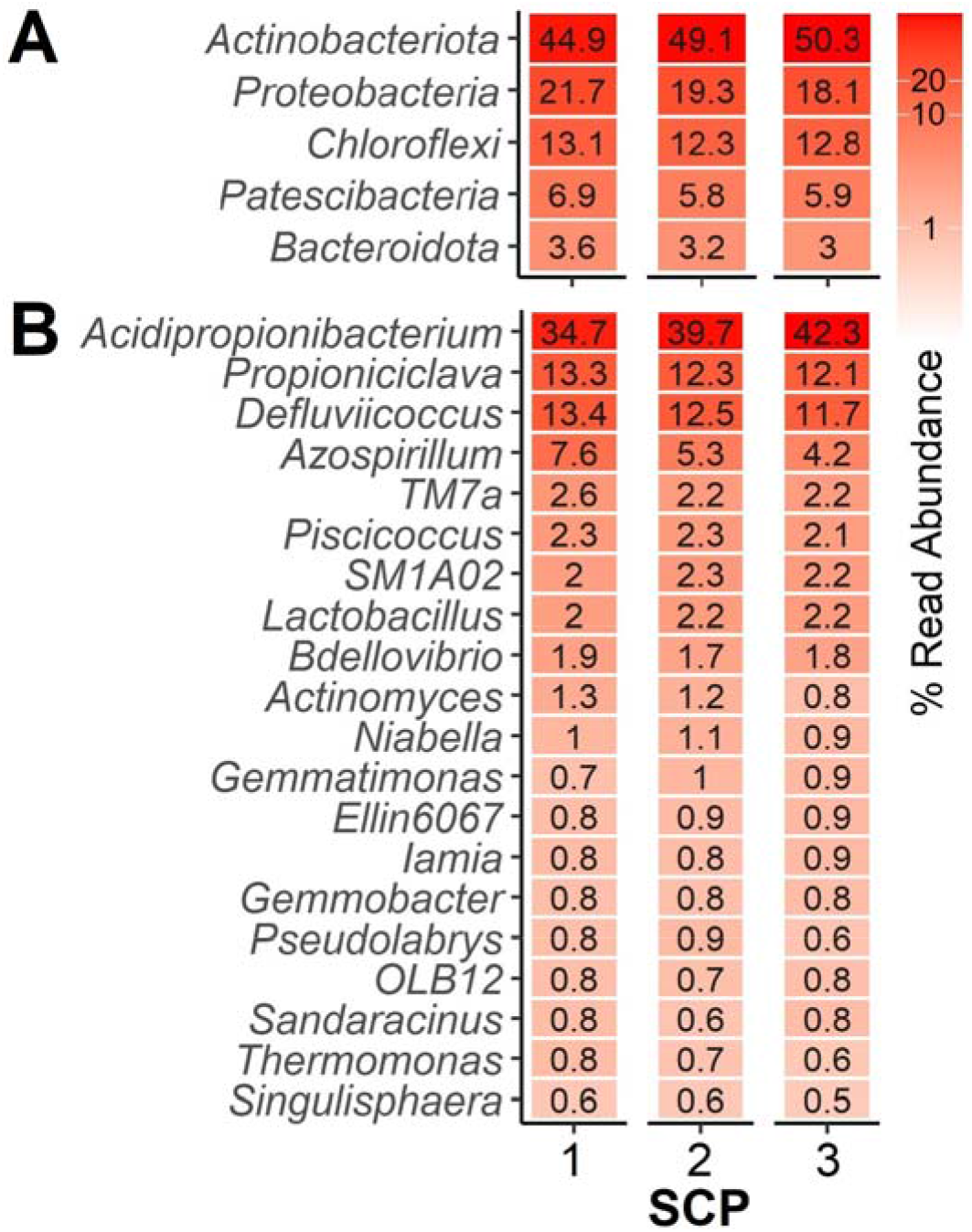
Microbial characterization of the microbial community-based SCP produced from soybean processing wastewaters, assessed through 16S rRNA gene amplicon sequencing. (**A**) Top 5 phyla and (**B**) top 20 genera, from three separate samples of the SCP mixture used in the study.

Based on the results of this preliminary trial, 50% of the fishmeal protein might be replaced with microbial community-based SCP meal without negatively affecting Asian seabass growth or survival in the short term. However, further studies are warranted for longer time effects and different protein replacement levels. Moreover, other fish species and sources of FPWW should also be explored. It has been shown that microbial community-based single SCP grown on wastewaters from potato and starch, brewery or dairy industries, meets several of the essential amino acid requirements of fish and shrimp (Vethathirri et al., 2021). However, FPWW of variable chemical composition may also lead to different mixed SCP communities (Vethathirri et al., 2022), thus more research is needed for a broader validation of this approach.

## 4. Conclusions

This preliminary trial study showed that 50% of the fishmeal protein might be replaced with microbial community-based SCP from soybean processing wastewater without affecting Asian seabass growth or survival, and that an SCP replacement diet may also lead less variable fish growth than with traditional fishmeal. Future studies should consider longer growth periods and higher fishmeal replacement levels, as well as additional aquacultured species and FPWW. Overall, we demonstrated that microbial community-based SCP has potential as an alternative value-added ingredient for aquaculture feed, which can help in the transition to a circular bioeconomy.

## Data availability

DNA sequencing data will be available at NCBI BioProjects PRJNA890376 upon publication.

## Author Contributions

ES, FY, DCPS and SW conceived the study and designed the experiment. SW and DCPS obtained the funding for the study. RSV and ES produced the microbial protein. FY performed the aquaculture trial. RSV and SST did the laboratory chemical analyses during SCP production. HYH and ES performed the molecular work. ES did the bioinformatics analyses. ES and FY interpreted the data, generated the results, and wrote the first draft of the manuscript. All authors reviewed the manuscript.

## Competing interests

The authors declare no competing interests.

## Funding Statement

This research was supported by the Singapore National Research Foundation (NRF) and Ministry of Education under the Research Centre of Excellence Program, and the NRF Competitive Research Programme [NRF-CRP21-2018-0006] “Recovery and microbial synthesis of high-value aquaculture feed additives from food-processing wastewater”.

## Notes

### Competing Interest Statement

The authors have declared no competing interest.

